# Post-infection treatment with a protease inhibitor increases survival of mice with a fatal SARS-CoV-2 infection

**DOI:** 10.1101/2021.02.05.429937

**Authors:** Chamandi S. Dampalla, Jian Zhang, Krishani Dinali Perera, Lok-Yin Roy Wong, David K. Meyerholz, Harry Nhat Nguyen, Maithri M. Kashipathy, Kevin P. Battaile, Scott Lovell, Yunjeong Kim, Stanley Perlman, William C. Groutas, Kyeong-Ok Chang

## Abstract

Severe acute respiratory syndrome coronavirus 2 (SARS-CoV-2) infection continues to be a serious global public health threat. The 3C-like protease (3CLpro) is a virus protease encoded by SARS-CoV-2, which is essential for virus replication. We have previously reported a series of small molecule 3CLpro inhibitors effective for inhibiting replication of human coronaviruses including SARS-CoV-2 in cell culture and in animal models. Here we generated a series of deuterated variants of a 3CLpro inhibitor, GC376, and evaluated the antiviral effect against SARS-CoV-2. The deuterated GC376 displayed potent inhibitory activity against SARS-CoV-2 in the enzyme and the cell-based assays. The K18-hACE2 mice develop mild to lethal infection commensurate with SARS-CoV-2 challenge doses and was proposed as a model for efficacy testing of antiviral agents. We treated lethally infected mice with a deuterated derivative of GC376. Treatment of K18-hACE2 mice at 24 hr post infection with a derivative (compound 2) resulted in increased survival of mice compared to vehicle-treated mice. Lung virus titers were decreased, and histopathological changes were ameliorated in compound 2-treated mice compared to vehicle-treated mice. Structural investigation using high-resolution crystallography illuminated binding interactions of 3CLpro of SARS-CoV-2 and SARS-CoV with deuterated variants of GC376. Taken together, deuterated GC376 variants have excellent potential as antiviral agents against SARS-CoV-2.

## INTRODUCTION

Coronaviruses are a large group of viruses that can cause a wide variety of diseases in humans and animals(1). They are single-stranded, positive-sense RNA viruses that belong to four genera, designated α, β, γ and δ coronaviruses, in the *Coronaviridae* family(2). Human coronaviruses (229E, NL63, OC43, and HKU1) generally cause mild upper respiratory infections. However, global outbreaks of new human coronavirus infections with severe respiratory disease have periodically emerged from animals, including Severe Acute Respiratory Syndrome Coronavirus (SARS-CoV)(3), Middle East Respiratory Syndrome Coronavirus (MERS-CoV)(4) and, most recently, SARS-CoV-2, the causative agent of COVID-19(5, 6). SARS-CoV-2 emerged in China in December 2019 and subsequently spread throughout the world. Ominously, the diversity of coronavirus strains in potential animal reservoirs suggests that emerging and reemerging pathogenic coronaviruses will continue to pose a significant threat to public health(7, 8). Currently, vaccines using different platforms have been developed or under development, and two vaccines just became available in the US for COVID-19 licensed for emergency use with more others expected to be available soon. The specific therapeutic interventions that are currently licensed include a nucleoside analogue remdesivir (Veklury®), a combination of remdesivir and a JAK inhibitor baricitinib, and a cocktail of anti-SARS-CoV-2 monoclonal antibodies. These treatments may diminish disease progression, but such effects are not found in most studies, indicating the urgent necessity to develop additional antiviral therapies(9–12).

The SARS-CoV-2 genome encodes two polyproteins which are processed by a 3C-like protease (3CLpro) and a papain-like protease. These viral proteases are essential for viral replication, making them attractive targets for drug development (13–18). It is furthermore acknowledged that, in addition to the development of effective vaccines, the concurrent identification of FDA-approved drugs that can be repurposed for use against SARS-CoV-2 may accelerate the development and implementation of effective countermeasures against the virus(19, 20). We previously described a series of 3CLpro inhibitors (including GC376) with activities against multiple coronaviruses, including SARS-CoV(21), MERS-CoV(13, 22) and SARS-CoV-2(13). GC376 was recently demonstrated in clinical trials to have efficacy against a fatal feline coronavirus infection, feline infectious peritonitis (FIP)(23, 24), and is currently in clinical development for treating FIP in cats. Mice expressing human angiotensin I-converting enzyme 2 (ACE2) receptor under the cytokeratin-18 (K18) promoter, designated as K18-hACE2 mice, were previously proposed as a model for efficacy testing of antiviral agents(25, 26). We report herein the results of our studies related to the synthesis and evaluation of deuterated GC376 variants which have enhanced antiviral activity and display efficacy in a fatal mouse model (K18-hACE2 mice) of SARS-CoV-2.

## RESULTS

### Deuterated variants of GC376 display potent inhibitory activity against SARS-CoV-2 in the enzyme and the cell-based assays

We synthesized deuterated variants based on GC376 (Figure S1, and Supporting information) and compared their inhibitory activities against SARS-CoV-2 to non-deuterated GC376 in the enzyme and the cell-based assays (Table 1). Three different variants of deuterated aldehyde compounds (compounds ***1*, **6**** and ***9*** with R1, R2 and R_3_, respectively) as well as their bisulfite adducts (compounds ***2*, **7**** and ***10***) were prepared for the testing. In addition, an α-ketoamide (compound ***5***) based on compound ***1*** and prodrug variations (compounds ***3***, ***4*, **8**** and ***11***) of the bisulfite adducts of aldehydes (compounds ***1***, ***6*** and ***9***) were synthesized for the testing. In the enzyme assay, the bisulfite adducts showed similar 50% inhibitory concentration (IC_50_) values as their aldehyde counterparts (Table 1C). The α-ketoamide derivative (compound ***5***) of compound ***1*** had markedly decreased potency in the enzyme assay. Likewise, prodrug counterparts (compounds ***3*, **4**, **8**** and ***11***) have significantly increased IC_50_ values compared to their aldehyde and bisulfite precursors (Table 1C). The deuterated compounds that were more effective than GC376 in the enzyme assay were tested in the cell-based assay. The 50% effective concentration (EC_50_) values of the tested deuterated compounds (compounds ***1***, ***2***, ***6*** and ***7***) (0.068 to 0.086 μM) were lower than GC376 by 2.67~3.38-fold in Vero E6 cells. All compounds, including GC376, did not show any cytotoxicity up to 100 μM (Table 1C).

**Table 1.** Structures and inhibitory activities of deuterated variants of GC376 against SARS-CoV-2 in the enzyme and cell-based assays. (A) R1, R2 and R3 deuterated moieties. (B) Structures of deuterated variants. (C) The activity and cytotoxicity of deuterated variants of GC376. IC_50_, the 50% inhibitory concentration determined in the enzyme assay; EC_50_, the 50% effective concentration determined in Vero E6 cells; and CC_50_, the 50% cytotoxic concentration determined in Vero E6 and CRFK cells. The values indicate the means and the standard deviations of the means.

| Compound | SARS-CoV-2 |  |  |
| --- | --- | --- | --- |
|  | IC<sub>50</sub> (μM) | EC<sub>50</sub> (μM) | CC<sub>50</sub> (μM) |
| **1** | 0.17±0.05 | 0.068±0.01 | > 100 |
| **2** | 0.18±0.04 | 0.086±0.02 | > 100 |
| **3** | 0.71±0.06 | N/D* | > 100 |
| **4** | 0.80±0.1 | N/D | > 100 |
| **5** | 1.31±0.55 | N/D | > 100 |
| **6** | 0.16±0.03 | 0.077±0.01 | > 100 |
| **7** | 0.19±0.04 | 0.079±0.01 | > 100 |
| **8** | 0.68±0.35 | N/D | > 100 |
| **9** | 0.24±0.01 | N/D | > 100 |
| **10** | 0.21±0.02 | N/D | > 100 |
| **11** | 0.77±0.12 | N/D | > 100 |
| **GC376** | 0.41±0.07 | 0.23±0.01 | > 100 |

\* N/D: not determined.

### Structures of 3CLpro of SARS-CoV and SARS-CoV-2 bound with deuterated variants of GC376

Compound ***2*** with a bisulfite adduct warhead and compound ***5*** with α-ketoamide were co-crystallized with the 3CLpro of SARS-CoV-2 and SARS-CoV and examined by X-ray crystallography. Examination of the active site of SARS-CoV-2 3CLpro revealed the presence of prominent difference electron density consistent with compound ***2*** covalently bound to the Sγ atom of Cys in each subunit (Figure 1A and B). Interestingly, the electron density was most consistent with the S-enantiomer at the newly formed stereocenter. Although the electron density in subunit B did contain a small “bulge” that may be due to the R-enantiomer, only one configuration was modeled. Compound ***2*** adopts the same binding mode in each subunit and forms identical hydrogen bond interactions with residues Phe^140^, His^163^, His^164^, Glu^166^ and Gln^189^ (Figure 1D and E). As we generally observed in studies of SARS-CoV 3CLpro, the electron density map was consistent with both the R and S-enantiomers of compound ***2*** at the new stereocenter formed by covalent attachment of the Sγ atom of Cys^145^ in the cocrystal structure of SARS-CoV 3CLpro (Figure 1C). Overall, the hydrogen bond interactions are nearly identical relative to SARS-CoV-2 3CLpro. The main difference is that a hydrogen bond is formed between His^41^ and the hydroxyl of compound ***2*** in the R-enantiomer and a long contact (3.29 Å) to the backbone N-atom of Ser^144^ with the hydroxyl of the S-enantiomer (Figure 1F). Notably, the hydroxyl in compound ***2*** bound to SARS-CoV-2 3CLpro is 3.38 Å and 3.39 Å from the N-atom of Ser^144^, which would be a weak hydrogen bond contact. The benzyl ring in both structures is positioned outward from the hydrophobic S_4_ subsite and are directed towards the surface as shown in Figure 1G, H and I. Notably, the structures of SARS-CoV-2 3CLpro in complex with nondeuterated G376 and its precursor aldehyde GC373 (PDB 6WTJ and 6WTK, respectively) adopts the same binding mode as that observed for compound ***2*** (Figure S2). Superposition yielded root-mean-square deviation (RMSD) deviations of 0.59 Å (GC376) and 0.55 Å (GC373) between Cα atoms for 299 residues aligned(27).

**Figure 1.**
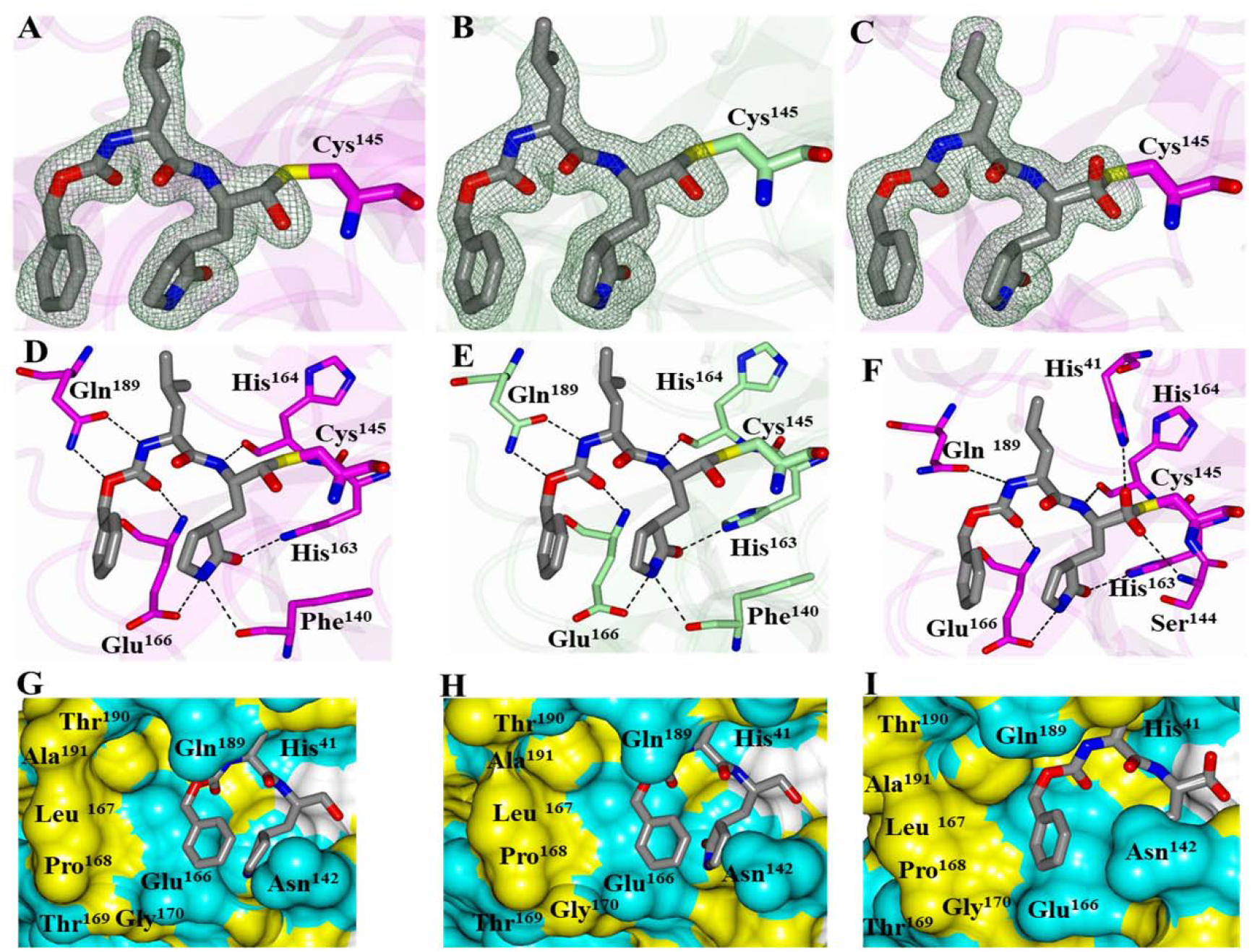
Cocrystal structures of SARS-CoV-2 3CLpro (**A**, **D**, **G: subunit A and B, E, H: subunit B**) and SARS-CoV 3CLpro (**C, F, I**) in complex with compound ***2***. Panels A-C show *F_o_-F_c_* omit maps (green mesh) contoured at 3σ. Panels D-F show hydrogen bond interactions (dashed lines) between the inhibitor and the 3CL protease. Panels G-I show electrostatic surface representation of the binding pocket occupied by the inhibitor. Neighboring residues are colored yellow (nonpolar), cyan (polar), and white (weakly polar).

The structures of SARS-CoV and SARS-CoV-2 3CLpro in complex with compound ***5*** also contained prominent difference in electron density consistent with the inhibitor covalently bound to the Sγ atom of Cys (Figure S3A and D). The entire inhibitor could be modeled in subunit A but was partially disordered in subunit B and the benzyl group in the S_4_ subsite could not be modeled for SARS-CoV. The inhibitor forms direct hydrogen bond interactions similar to compound ***2*** as shown in Figure S3B and E. The benzyl group in subunit A of both SARS-CoV and SARS-CoV-2 3CLpro is positioned near hydrophobic residues within the S_4_ subsite (Figure S3C and F). The benzyl ring in the α-ketoamide region of the inhibitor is positioned near a cleft formed by Asn^142^/Gly^143^ in both structures. Interestingly, the compound bound to subunit B of SARS-CoV-2 3CLpro adopts a conformation similar to that observed for compound ***2*** in which the benzyl group is directed away from the S_4_ subsite and towards the surface (Figure S4). Therefore, it appears the structure of SARS-CoV-2 3CLpro in complex with compound ***5*** serendipitously contains two binding modes of the inhibitor.

### Treatment with compound *2* at 24 hrs after SARS-CoV-2 infection demonstrates efficacy against fatal SARS-CoV-2 infection in K18-hACE2 mice

Compound ***2*** was tested in SARS-CoV-2-infected K18-hACE2 mice for protective efficacy, because it potently inhibited SARS-CoV-2 in the cell-based assay described above. The dose curve of compound ***2*** against SARS-CoV-2 in cell culture is shown in Figure 2A. In the first experiment, infection with 2×10^3^ pfu per mouse led to body weight loss in all vehicle-treated mice resulting in 50% survival by 9 dpi (Figure 2B). Mice treated with compound ***2*** (100 mg/kg/day, once a day) starting from 24 hr post infection (1 dpi) lost body weight, but loss was less severe compared to vehicle-treated mice with statistically significant differences (*0.002<p<*0.049) on most days between 5-10 dpi (Figure 1C). All compound ***2***-treated mice gradually gained body weight and were alive at the end of the study (15 dpi) (Figure 1C), although survival of these mice was not statistically different *(p=0.18)* compared to vehicle-treated mice, likely due to lower fatality of control mice. In the second experiment, a higher virus challenge dose (5×10^3^ pfu per mouse) was given before treatment with vehicle or compound ***2*** (125 mg/kg/day, once a day) started 24 hr post infection. The vehicle-treated mice exhibited greater body weight loss than those with the lower virus challenge (experiment 1), and none of the mice (N=4) survived past 8 dpi (Figure 1D). Weight losses of mice treated with compound ***2*** were significantly less than those with vehicle treatment at 3, 5, and 6 dpi (0.009<*p<0.017*), and compound ***2***-treated mice started to gain weight from 10 dpi (Figure 1E). In contrast to 0% survival in vehicle-treated mice, 5 out of 6 compound ***2***-treated mice (83%) survived at the end of the study (15 dpi), resulting in significant improved survival of compound ***2***-treated mice *(P = 0.011).*

**Figure 2.**
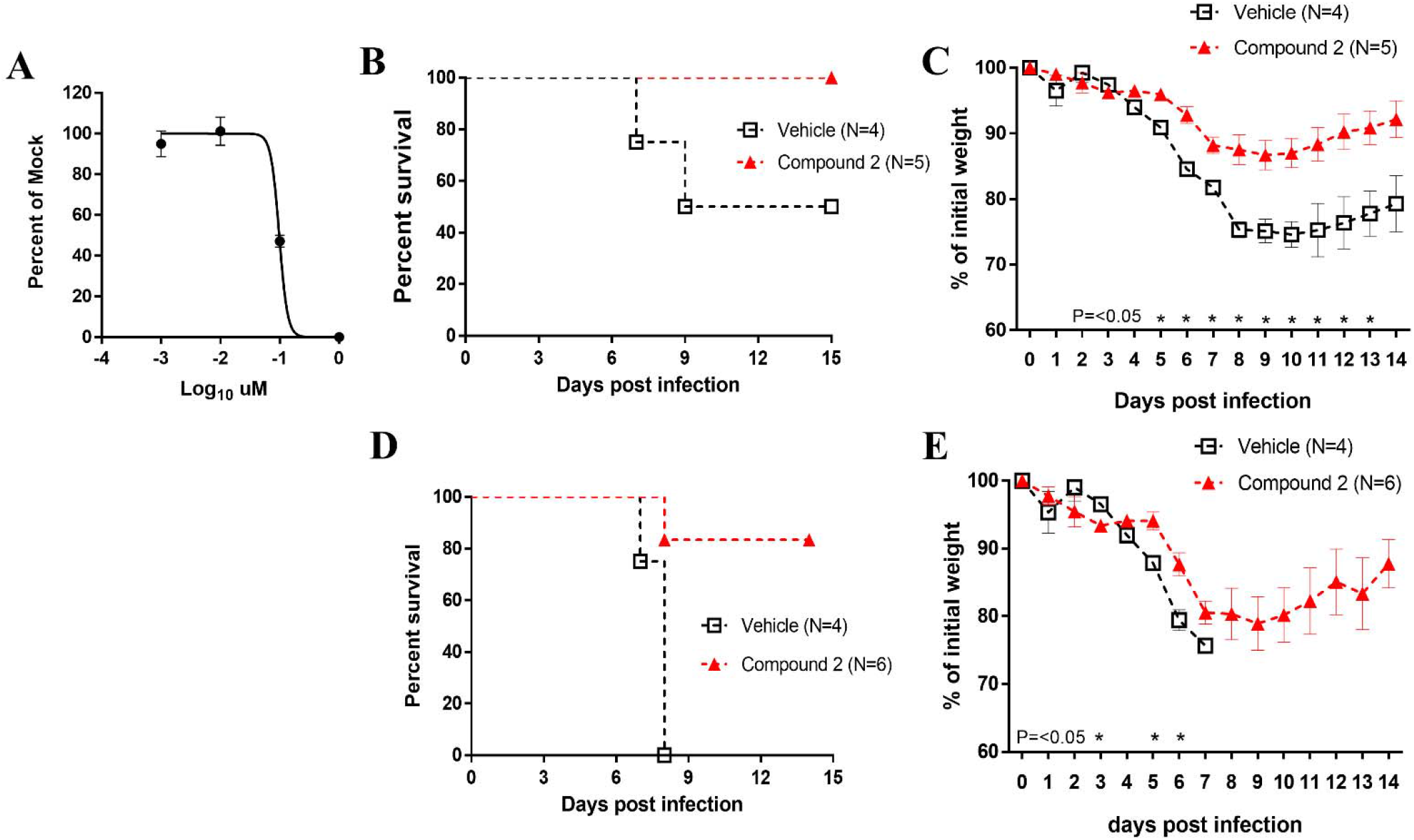
Therapeutic treatment of K18-hACE2 mice infected with SARS-CoV-2. (A) A dose-dependent curve for compound ***2*** against SARS-CoV-2 in cell culture. Confluent Vero E6 cells were inoculated with SARS-CoV-2, and medium containing various concentrations of compound ***2*** and agar was applied to the cells. After 48-72 hr, plaques in each well were counted, and EC_50_ values were determined by GraphPad Prism software. (B and C) The K18-hACE2 mice infected with SARS-CoV-2 with 2×10^3^ plaque forming unit (pfu)/mouse were treated with compounds ***2*** at 100 mg/kg once per day (N=5) or vehicle (N=4) starting at 1-day post-infection (dpi) for up to 10 days, and survival (B) and weight (C) were monitored for 15 days. (D and E) The K18-hACE2 mice infected with SARS-CoV-2 with 5 x10^3^ pfu/mouse were treated with compound ***2*** at 125 mg/kg once per day (N=6) or vehicle (N=4) starting at 1 dpi for up to 10 days, and survival (D) and weight (E) were monitored for 15 days. The data points represent the means and the standard deviations of the means. The analysis of survival curves between groups was performed using a Log-rank (Mantel-Cox) test in GraphPad Prism software. The symbols and the bars in Panels C and E represent the means and the standard deviations of the means. Asterisks indicate statistical difference between vehicle and compound ***2***-treated groups determined using multiple T-test in GraphPad Prism software *(p<0.05)*.

### Treating infected K18 hACE2 mice with compound *2* reduces viral titers and histopathological changes in the lungs

Mice were infected with 5×10^3^ pfu SARS-CoV-2 virus and treated with compound ***2*** or vehicle starting from 1 dpi. Lung virus titers peaked at 2 dpi and decreased at 5 dpi in both groups. Virus titers were statistically lower in compound ***2***-treated mice compared to vehicle-treated mice at 5 dpi by approximately 10-fold *(P=0.0028)* (Figure 3A). Lung pathology in vehicle-treated mice included diffuse alveolar damage with progressive alveolar or interstitial lesions characterized by edema, inflammation and focal cytomegaly in some alveolar lining cells. Additional features include an accumulation of immune effector cells, including granulocytes and macrophages, evidence of cell death, hemorrhage, hyaline membranes, and occasional vascular thrombi. Histopathological observations were in agreement with improved survival observed in SARS-CoV-2 infected animals treated with compound ***2*** (Figure 3B). At 2 dpi, alveolar edema was seen in some lung tissue sections (average score 0.5) in the lungs from animals treated with vehicle (Figure 3B a and b), but there were few lesions in the lungs from compound ***2***-treated animals (average score 0) (Figure 3B c and d). Mild perivascular infiltrates were seen in lungs from both vehicle and compound ***2***-treated animals at 2 dpi (Figure 3B a-d). At 5 dpi, while severe edema (average score 4) and perivascular infiltrates (average score 3.5) were evident in the lungs from vehicle-treated animals (Figure 3B e and f), mild edema (average score 2) and perivascular infiltrations (average score 2) were observed in the lungs of compound ***2***-treated animals (Figure 3B g and h). K18-hACE2 mice infected with SARS-coV-2 sometimes develop encephalitis. Low levels of virus but no pathological changes were detected in vehicle-treated mice (Fig 3C). In contrast, neither pathological changes nor infectious virus was detected at 2 and 5 dpi in the brains of mice treated with compound ***2*** (Fig 3C)

**Figure 3.**
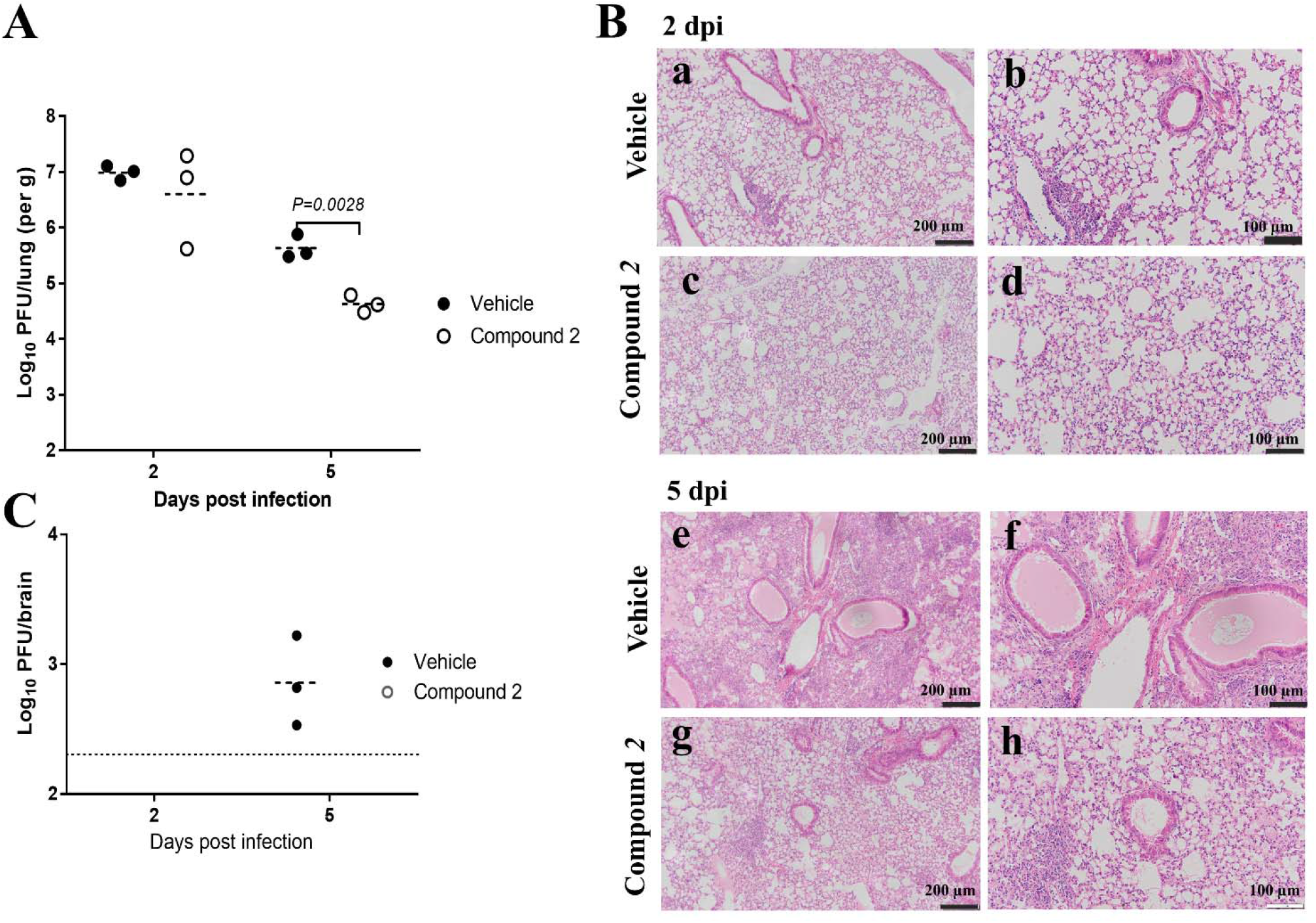
Lung virus titers and histopathology of K18-hACE2 mice infected with SARS-CoV-2 and treated with vehicle or compound ***2*** starting at 1-day post infection (dpi). The lungs and the brains were collected at 2 and 5 dpi for virus titration (A and C) and histopathology (B). (A) Lung virus load in vehicle- or compound ***2***-treated groups. Each symbol represents individual mouse, and the dashed line represents the means while the dotted line is the limit of detection (200 plaque forming unit, pfu). Confluent Vero E6 cells were inoculated with serial dilutions of lung homogenates and agar was applied to the cells. After 48-72 hr, plaques in each well were counted and pfu per gram was calculated. Statistical significance was determined using multiple T-test in GraphPad Prism software *(p<0.05).* (B) Lungs were examined for edema and for hyaline membrane formation. Lung sections were stained with hematoxylin and eosin for histopathology at 2 (a to d) or 5 dpi (e to h). Histopathology images are shown at either 10X (a, c, e and g) with a 200 μm scale bar or 20X (b, d, f and h) with a 100 μm scale bar for vehicle control (a, b, e and f) and compound ***2***-treat groups (c, d, g and h). (C) Brain virus load in vehicle- or compound ***2***-treated groups. Virus titration and statistical analysis were performed following the same procedures described in for the lung samples (A).

## DISCUSSION

The advent of SARS-CoV-2, the causative agent of COVID-19, has provided the impetus behind worldwide efforts to develop effective countermeasures against the virus for the treatment of COVID-19, including the use of repurposed drugs [reviews(19, 20)]. Indeed, remdesivir, a nucleoside analogue, which was originally developed and FDA-approved for treating Ebola virus infection has been shown to be a potent inhibitor of SARS-CoV-2(28-30) and recently approved for COVID-19. However, effects in patients have been modest, with some studies showing no efficacy(9-12). Multiple FDA-approved drugs which exert their antiviral effects by impeding key steps in the viral lifecycle, including virus entry and fusion, and viral replication, among others, are currently under intense investigation for use against SARS-CoV-2(19, 20, 31–33). Efforts in developing small molecule inhibitors targeting the virus proteases of SARS-CoV- 2 have focused on blocking 3CLpro(17, 34, 35). Recently, GC376, a 3CL protease inhibitor under commercial development for FIP, was reported to have anti-SARS-CoV-2 activity by us(13) and other groups(27, 36, 37), which suggests this compound is a lead compound for COVID-19 amenable to further optimization. Based on the multiple potential advantages frequently accrued from the introduction of deuterium in a drug, such as improved pharmacokinetics, reduction in toxicity, and enhanced potency(38-40), it was envisaged that deuterated variants of GC376 could function as therapeutics with superior characteristics compared to the corresponding non-deuterated GC376 drug. Therefore, we generated three deuterated variants of GC376 by replacing hydrogen with deuterium at the metabolic soft spots encompassed in the R site (aromatic ring and benzylic carbon) and evaluated their activity in the enzyme and the cell-based assays. All three R site-deuterated variants (aldehydes and their bisulfite adducts) showed modestly increased potency compared to GC376, which was more apparent in the cell-based assays than in the enzyme assay (Table 1C).

Crystal structures of deuterated GC376 (compound ***2***) and the 3CLpro of SARS-CoV-2 revealed that deuteration did not alter the interactions between GC376 and 3CLpro which are reported by other groups(27, 37, 41). It may be speculated that the enhanced activity of the deuterated compounds can be attributed to tighter binding to the target, which was observed with other deuterated compounds(38) or improved physicochemical properties of the compound. However, further study is needed to understand the mechanism. Substitution of the aldehyde warhead in compound ***1*** with an α-ketoamide (compound ***5***) significantly decreased potency, which confirms earlier findings by us(21, 42) that ketoamide is less suitable for coronavirus 3CLpro inhibition. Employing an OCOmethyl or n-pentyl ester derivative of bisulfite adducts to produce prodrug variants (compounds ***3***, ***4*, **8**** and ***11***) led to reduced potency in the enzyme assay, which may be due to inefficient conversion to the active compound in the enzyme assay.

Most animals (hamsters, ferrets, and non-human primates) experimentally infected with SARS-CoV-2 serve as good models for asymptomatic, mild, and moderate SARS-CoV-2 infection and for studies of viral transmission. However, there are limited animal models that recapitulate the key features of severe pathogenesis in humans with COVID-19. K18-hACE2 mice infected with SARS-CoV-2 develop a mild to lethal infection dependent upon virus challenge doses and was previously proposed as a model for efficacy testing of antiviral agents(26, 43, 44). Mice with fatal virus infection showed viral replication in the lungs with inflammation and virus-induced histopathological changes that resemble severe COVID-19 infection in humans. In this model, pre- and postinfection treatment efficacy of human convalescent plasma (CP) from a recovered COVID-19 patient was previously studied(26). In the current study, once daily administration of compound ***2*** starting at 24 hr post infection to mice infected with 50% or 100% lethality resulted in significant reductions in body weight loss and nearly complete survival. The kinetics of virus clearance was enhanced, and lung pathological changes were diminished by drug treatment.

In summary, we show that deuterated variants of GC376 are potent inhibitors of SARS-CoV-2 replication and significantly enhance survival of infected mice. Importantly, this approach has wide applicability, and strategic deuteration can be extended to multiple FDA-approved drugs currently under investigation as COVID-19 therapeutics with potential improvement in clinical outcomes.

## Materials and Methods

### Study Design

The primary objective of this study was to evaluate the antiviral activity of deuterated variants of GC376, a protease inhibitor that is currently under investigation for the treatment of feline infectious peritonitis and COVID-19, against SARS-CoV-2 *in vitro,* as well as antiviral efficacy in a fatal mouse model of SARS-CoV-2 infection.

### Biocontainment and biosafety of coronaviruses

All studies with SARS-CoV-2 were performed in biosafety level 3 facilities at the University of Iowa. All experiments were conducted under protocols approved by the Institutional Biosafety Committee at the University of Iowa according to guidelines set by the Biosafety in Microbiological and Biomedical Laboratories, the U.S. Department of Health and Human Services, the U.S. Public Health Service, the U.S. Centers for Disease Control and Prevention, and the National Institutes of Health.

### Synthesis of deuterated variants of GC376 3CLpro inhibitors

Compounds ***1-11*** were readily synthesized using a reaction sequence similar to the one previously reported by us (Figure S1)(45-47), and are listed in Table 1B. Briefly, the deuterated alcohol inputs (Table 1A) were reacted with (L) leucine isocyanate methyl ester to yield the corresponding dipeptidyl methyl esters which were then hydrolyzed to the corresponding acids with lithium hydroxide in aqueous tetrahydrofuran. Subsequent carbonyl dimidazole (CDI)-mediated coupling of the acids to glutamine surrogate methyl ester(48) furnished the dipeptidyl methyl esters. Lithium borohydride reduction yielded the alcohols which were then oxidized to the corresponding aldehydes with Dess-Martin periodinane reagent. The bisulfite adducts were generated by treatment with sodium bisulfite in aqueous ethanol and ethyl acetate(49). An alternative convergent synthesis of compounds ***1-11*** entailed the activation of a deuterated alcohol with disuccinimidoyl carbonate followed by sequential coupling with a dipeptidyl amino alcohol and oxidation with Dess-Martin periodinane. The synthesis of α-ketoamide compound **5** was accomplished by reacting the Z-protected dipeptidyl aldehyde with benzyl isonitrile to yield the α-hydroxyketoamide followed by Dess-Martin oxidation. Prodrug compounds ***3***, ***4***, ***8*** and ***11*** were synthesized by refluxing the aldehyde bisulfite adduct with acetic anhydride or n-hexanoic anhydride(47).

### Fluorescence resonance energy transfer (FRET) enzyme assay

The cloning, expression, and purification of the 3CLpro of SARS-CoV-2 were conducted by a standard method described previously by our lab(13) and others(15, 17, 50). The codon-optimized cDNA of full length 3CLpro of SARS-CoV-2 (GenBank number MN908947.3) fused with sequences encoding 6 histidine at the N-terminal was synthesized by Integrated DNA Technologies (Coralville, IA). The synthesized gene was subcloned into the pET-28a (+) vector. The FRET enzyme assays were conducted as described previously(17). Briefly, stock solutions of compounds ***1-11*** were prepared in DMSO and diluted in assay buffer which was comprised of 20 mM HEPES buffer, pH 8, containing NaCl (200 mM), EDTA (0.4 mM), glycerol (60%), and 6 mM dithiothreitol (DTT). The SARS-CoV-2 3CL protease was mixed with serial dilutions of each compound or with DMSO in 25 μL of assay buffer and incubated at room temperature for 1 hr (SARS-CoV-2 and SARS-CoV), followed by the addition of 25 μL of assay buffer containing substrate (FAM-SAVLQ/SG-QXL®520, AnaSpec, Fremont, CA). The substrate was derived from the cleavage sites on the viral polyproteins of SARS-CoV. Fluorescence readings were obtained using an excitation wavelength of 480 nm and an emission wavelength of 520 nm on a fluorescence microplate reader (FLx800; Biotec, Winoosk, VT) at 1 hr following the addition of the substrate. Relative fluorescence units (RFU) were determined by subtracting background values (substrate-containing well without protease) from the raw fluorescence values, as described previously(13). The dose-dependent FRET inhibition curves were fitted with a variable slope using GraphPad Prism software (GraphPad, La Jolla, CA) to determine the 50% inhibitory concentration (IC50) values of the compounds, which are listed in Table 1C.

### Cell-based assay for antiviral activity

Compounds ***1***, ***2***, ***6*** and ***7*** were investigated for their antiviral activity against the replication of SARS-CoV-2. Briefly, confluent Vero E6 cells were inoculated with SARS-CoV-2 at 50-100 plaque forming units/well, and medium containing various concentrations of each compound and agar was applied to the cells. After 48-72 hr, plaques in each well were counted. The 50% effective concentration (EC_50_) values were determined by GraphPad Prism software using a variable slope (GraphPad, La Jolla, CA)(17).

### Nonspecific cytotoxic effects

The 50% cytotoxic concentrations (CC_50_) of compounds ***1-11*** were determined in Vero E6 and CRFK cells. Confluent cells grown in 96-well plates were incubated with various concentrations (1 to 100 μM) of each compound for 72 hr. Cell cytotoxicity was measured by a CytoTox 96 nonradioactive cytotoxicity assay kit (Promega, Madison, WI), and the CC_50_ values were calculated using a variable slope by GraphPad Prism software (Table 1C).

### X-ray crystallographic studies: protein purification, crystallization, and data collection

Purified SARS-CoV-2 3CLpro and SARS-CoV 3CLpro were concentrated to 22.0 mg/mL (0.64 mM) and 9.6 mg/mL (0.28 mM), respectively, in 100mM NaCl, 20mM Tris pH 8.0 for crystallization screening. All crystallization experiments were set up using an NT8 drop-setting robot (Formulatrix Inc.) and UVXPO MRC (Molecular Dimensions) sitting drop vapor diffusion plates at 18 °C. 100 nL of protein and 100 nL crystallization solution were dispensed and equilibrated against 50 μL of the latter. Stock solutions of compounds ***2*** and ***5*** (100 mM) were prepared in DMSO and the 3CLpro:inhibitor complexes were prepared by adding 2 mM ligand to the proteases and incubating on ice for 1 hr. Crystals were obtained in 1-2 days from the following conditions. **SARS-CoV 3CLpro complex with compound *2*:** Index HT screen (Hampton Research) condition H4 (30% (w/v) PEG 3350, 200 mM ammonium citrate pH 7.0). **SARS-CoV 3CLpro complex with compound *5*:** Berkeley screen (Rigaku Reagents) condition B1 (30% (w/v) PEG 3350, 100 mM Tris pH 8.5, 400 mM sodium chloride). **SARS-CoV-2 3CLpro complex with compound *2*:** Index HT screen (Rigaku Reagents) condition G8 (25% (w/v) PEG 3350, 100 mM Hepes pH 7.5, 200 mM ammonium acetate). **SARS-CoV-2 3CLpro complex with compound *5*:** Index HT condition D11 (28% PEG 2000 MME, 100 mM Bis-Tris pH 6.5). Samples were transferred to a fresh drop composed of 80% crystallization solution and 20% (v/v) PEG 200 and stored in liquid nitrogen. All X-ray diffraction data were collected using a Dectris Eiger2 X 9M pixel array detector at the Advanced Photon Source IMCA-CAT beamline 17-ID except for the data for the SARS-CoV-2 3CLpro complex with compound ***5*** which were collected at the National Synchrotron Light Source II (NSLS-II) AMX beamline 17-ID-1.

### Structure solution and refinement

Intensities were integrated using XDS via Autoproc(51) and the Laue class analysis and data scaling were performed with Aimless(52). Structure solution was conducted by molecular replacement with Phaser(53) using previously determined structures of SARS-CoV 3CLpro (PDB 6W2A) and SARS-CoV-2 3CLpro (PDB 6XMK) as the search models **(13)**. Structure refinement and manual model building were conducted with Phenix(54) and Coot(55), respectively. Disordered side chains were truncated to the point for which electron density could be observed. Structure validation was conducted with Molprobity(56), and figures were prepared using the CCP4MG package(57). Superpositions were performed using GESAMT(58). Crystallographic data are provided in Table S1.

### Animal care and ethics statement

*In vivo* studies were performed in animal biosafety level 3 facilities at the University of Iowa. All experiments were conducted under protocols approved by the Institutional Animal Care and Use Committee at the University of Iowa according to guidelines set by the Association for the Assessment and Accreditation of Laboratory Animal Care and the U.S. Department of Agriculture.

### Post-infection treatment in a mouse model of SARS-CoV-2 infection

Compound ***2*** was examined for efficacy using 7-8 week-old female K18-hACE2 mice infected with SARS-CoV-2(26). In the first experiment, animals were divided into two groups (N=4 for vehicle or N=5 for compound ***2***) and were lightly anesthetized with ketamine/xylazine prior to infection with 50 μl of 2×10^3^ pfu SARS-CoV-2 [The 2019n-CoV/USA-WA1/2019 strain of SARS-CoV-2 (accession number: MT985325.1)] via intranasal inoculation. Compound ***2*** was formulated in 10% ethanol and 90% PEG400 and given to mice from 1 (24 hr post infection) to 10 days post infection (dpi) at 100 mg/kg/day (once per day) via intraperitoneal administration. Control mice received vehicle. Animals were weighed daily and monitored for 15 days. Animals were euthanized when an animal lost 30% of initial weight or at 15 dpi. In the second experiment, mice were divided into two groups (N=4 for vehicle or N=6 for compound ***2***) and infected with 50 μl of 5×10^3^ pfu SARS-CoV-2 via intranasal inoculation. Compound ***2*** was given to mice from 1 (24 hr post infection) to 10 dpi at 125 mg/kg/day (once per day) via intraperitoneal administration. The third experiment was conducted simultaneously with the second one to determine virus titers and histopathology in the lungs of mice treated with vehicle or compound ***2*** (N=3 for virus titration and N=1 or 2 for histopathology for each group at each dpi). Animals were euthanized at 2 or 5 dpi, and lungs and brains were removed aseptically and disassociated with a manual homogenizer in 1X PBS. The homogenized lung and brain tissues were briefly centrifuged, and supernatants were removed. Virus titration was conducted in Vero E6 cells against SARS-CoV-2(26). For histopathology, lungs and brains were fixed with 10% formalin, and hematoxylin and eosin (HE) stained tissues were examined by a veterinary pathologist using the post-examination method of masking(26). Lung tissues were evaluated for edema (0-4) using distribution-based ordinal scoring: 0 - none; 1 - <25% of field; 26-50% of field; 51-75% field and >75% of field. Perivascular inflammation was evaluated by severity-based ordinal scoring: 0 - none; 1 - scant solitary cellular infiltrates that do not form aggregates; 2 - Mild infiltrates that form loose cuff (~1 cell thickness) around vessel; 3 - infiltrates form a distinct perivascular aggregate ~2-4 cells thick; and 4-large perivascular aggregates (>4 cells thick that extend to compress adjacent tissues.

### Statistical analysis

Multiple T-test were used to analyze body weight change and lung virus titers between groups using GraphPad Prism Software version 6 (San Diego, CA). Log-rank (Mantel-Cox) test was used for analysis of survival curves between groups using GraphPad Prism. *P* < 0.05 was considered statistically significant.

## Acknowledgments

We thank David George for the technical support. This work is supported in part by grants from the National Institutes of Health (R01 AI109039 to K.O.C. and P01 AI060699 and R01 AI129269 to S. P.). Use of the IMCA-CAT beamline 17-ID at the Advanced Photon Source was supported by the companies of the Industrial Macromolecular Crystallography Association through a contract with Hauptman-Woodward Medical Research Institute. This research used resources of the Advanced Photon Source, a U.S. Department of Energy (DOE) Office of Science User Facility, operated for the DOE Office of Science by Argonne National Laboratory under Contract No. DE-AC02-06CH11357. Extraordinary facility operations were supported in part by the DOE Office of Science through the National Virtual Biotechnology Laboratory, a consortium of DOE national laboratories focused on the response to COVID-19, with funding provided by the Coronavirus CARES Act. This research used the AMX beamline of the National Synchrotron Light Source II, a U.S. Department of Energy (DOE) Office of Science User Facility operated for the DOE Office of Science by Brookhaven National Laboratory under Contract No. DE-SC0012704. The Center for BioMolecular Structure (CBMS) is primarily supported by the National Institutes of Health, National Institute of General Medical Sciences (NIGMS) through a Center Core P30 Grant (P30GM133893), and by the DOE Office of Biological and Environmental Research (KP1605010).

## Supplementary Information

### Synthesis of deuterated 3CLpro inhibitors

#### General

Reagents and dry solvents were purchased from various chemical suppliers (Sigma-Aldrich, Acros Organics, Chem-Impex, TCI America, Oakwood chemical, APExBIO, Cambridge Isotopes and Fisher) and were used as obtained. Silica gel (230-450 mesh) used for flash chromatography was purchased from Sorbent Technologies (Atlanta, GA). Thin layer chromatography was performed using Analtech silica gel plates. Visualization was accomplished using UV light and/or iodine. NMR spectra were recorded in CDCl_3_ or DMSO-d_6_ using a Varian XL-400 spectrometer. Melting points were recorded on a Mel-Temp apparatus and are uncorrected. High resolution mass spectrometry (HRMS) was performed at the University of Missouri-Kansas City Mass Spectroscopy Core facility, using an LCMS-9030 Q-TOF mass spectrometer equipped with Nexera X3 (40-series) UHPLC system, dual ion source (DUIS) and quadrupole time of flight (Q-ToF) mass analyzer (Shimadzu, Columbia, MD). Purity was >95% as evidenced by proton NMR analysis.

#### Compound *1*

Phenylmethyl-d2 ((S)-4-methyl-1-oxo-1 -(((S)-1-oxo-3-((S)-2-oxopyrrolidin-3-yl)propan-2-yl)amino)pentan-2-yl)carbamate (***1***). Yield (57%), mp 45-48 ^0^C. ^1^H NMR (400 MHz, CDCl3): δ 9.48 (s, 1H), 8.32 (d, *J* = 5.9 Hz, 1H), 7.38 - 7.28 (m,5H), 6.18 (s, 1H), 5.44 (d, *J* = 8.6 Hz, 1H), 4.40 - 4.29 (m, 2H), 3.36 - 3.29 (m, 2H), 2.52 - 2.22 (m, 2H), 2.01 - 1.79 (m, 2H), 1.79 - 1.64 (m, 3H), 1.60 - 1.50 (m, 1H), 0.97 (d, *J* = 5.7 Hz, 6H). HRMS m/z: [M+H]^+^ Calculated for C_21_H_28_D_2_N_3_O_5_: 406.2311. Found: 406.2314; m/z: [M+Na]^+^ Calculated for C_21_H_28_D_2_N_3_NaO_5_: 428.2131. Found: 428.2136.

#### Compound *2*

Sodium (2S)-1-hydroxy-2-((S)-4-methyl-2-(((phenylmethoxy-d2)carbonyl)amino)pentanamido)-3-((S)-2-oxopyrrolidin-3-yl)propane-1-sulfonate (***2***). Yield (69%), mp 118-120 ^0^C. ^1^H NMR (400 MHz, DMSO-d_6_) δ 7.65 (d, *J* = 9.2 Hz, 1H), 7.59 (d, *J* = 9.3 Hz, 1H), 7.49 (d, *J* = 8.4 Hz, 1H), 7.33 - 7.27 (m, 5H), 5.49 (d, *J* = 6.2 Hz, 1H), 5.33 (d, *J* = 6.0 Hz, 1H), 4.27 - 4.19 (m, 1H), 3.88 - 3.83 (m, 1H), 3.18j - 2.96 (m, 2H), 2.21 - 1.89 (m, 2H), 1.67 - 1.50 (m, 2H), 1.50 - 1.38 (m, 4H), 0.83 (d, *J* = 2.9 Hz, 6H). HRMS m/z: [M+H]^+^ Calculated for C_21_H_29_D_2_N_3_NaO_8_S: 510.1855, Found: 510.1838.

#### Compound *3*

Sodium (5S,8S)-5-isobutyl-3,6,11-trioxo-8-(((S)-2-oxopyrrolidin-3-yl)methyl)-1-phenyl-2,10-dioxa-4,7-diazadodecane-9-sulfonate-1,1-d2 (***3***). Yield (72%), mp 110-113 ^0^C. ^1^H NMR (400 MHz, DMSO-dø) δ 7.70 - 7.62 (m, 2H), 7.49 (s, 1H), 7.41 - 7.30 (m, 5H), 5.10 (d, *J* = 8.9 Hz, 1H), 4.41 - 3.80 (m, 2H), 3.18 - 3.03 (m, 2H), 2.30 - 1.82 (m, 2H), 1.77 (s, 3H), 1.69 - 1.32 (m, 6H), 0.83 (d, *J* = 5.9 Hz, 6H). HRMS m/z: [M+H]^+^ Calculated for C_23_H_31_D_2_N_3_NaO_9_S: 552.1960. Found: 552.1953; m/z: [M+Na]^+^ Calculated for C_23_H_30_D_2_N_3_Na_2_O_9_S: 574.1780, Found: 574.1777. m/z: [M]^-^ Calculated for C_23_H_30_D_2_N_3_O_9_S: 528.1984, Found: 528.1993.

#### Compound *4*

Sodium (5S,8S)-5-isobutyl-3,6,11-trioxo-8-(((S)-2-oxopyrrolidin-3-yl)methyl)-1-phenyl-2,10-dioxa-4,7-diazahexadecane-9-sulfonate-1,1-d2 (***4***). Yield (64%), mp 100-104 ^0^C. ^1^H NMR (400 MHz, DMSO-d_6_) δ 7.68 (d, *J* = 10.4 Hz, 1H), 7.51 (d, *J* = 8.2 Hz, 1H), 7.47 (s, 1H), 7.42 - 7.23 (m, 5H), 5.11 (d, *J* = 8.7 Hz, 1H), 4.34 - 4.00 (m, 1H), 4.00 - 3.83 (m, 1H), 3.21 - 2.91 (m, 2H), 2.41 - 1.99 (m, 4H), 1.99 - 1.74 (m, 1H), 1.74 - 1.33 (m, 7H), 1.35 - 1.16 (m, 4H), 0.86 (d, *J* = 12.6 Hz, 9H). HRMS m/z: [M+H]^+^ Calculated for C_27_H_39_D_2_N_3_NaO_9_S: 608.2586. Found: 608.2581; m/z: [M+Na]^+^ Calculated for C_27_H_38_D_2_N_3_Na_2_O_9_S: 630.2406, Found: 630.2404; m/z: [M]^-^ Calculated for C_27_H_38_D_2_N_3_O_9_S: 584.2610, Found: 584.2623.

#### Compound *5*

Phenylmethyl-d2 ((S)-1-(((S)-4-(benzylamino)-3,4-dioxo-1-((S)-2-oxopyrrolidin-3-yl)butan-2-yl)amino)-4-methyl-1-oxopentan-2-yl)carbamate (***5***). Yield (40%), mp 124-128 ^0^C. ^1^H NMR (400 MHz, CDCl3) δ 8.43 (d, *J* = 6.1 Hz, 1H), 7.49 (t, *J* = 6.2 Hz, 1H), 7.37 - 7.19 (m, 10H), 6.89 (s, 1H), 5.73 (d, *J* = 8.7 Hz, 1H), 4.48 - 4.42 (m, 2H), 4.42 - 4.17 (m, 2H), 3.32 - 3.09 (m, 2H), 2.57 - 2.43 (m, 1H), 2.43 - 2.29 (m, 1H), 2.29 - 2.12 (m, 1H), 1.96 - 1.81 (m, 1H), 1.81 - 1.56 (m, 3H), 1.56 - 1.45 (m, 1H), 0.93 (d, *J* = 6.5 Hz, 6H). HRMS m/z: [M+H]^+^ Calculated for C_29_H_35_D_2_N_4_O_6_: 539.2838, Found: 539.2842, m/z: [M+Na]^+^ Calculated for C_29_H_34_D_2_N_4_NaO_6_: 561.2658, Found: 561.2661.

#### Compound *6*

(Phenyl-d5)methyl ((S)-4-methyl-1-oxo-1-(((S)-1-oxo-3-((S)-2-oxopyrrolidin-3-yl)propan-2-yl)amino)pentan-2-yl)carbamate (***6***). Yield (62%), mp 40-43 ^0^C. ^1^H NMR (400 MHz, DMSO-d_6_) δ 9.40 (s, 1H), 8.48 (d, *J* = 7.6 Hz, 1H), 7.49 (s, 1H), 7.38 (d, *J* = 8.3 Hz, 1H), 5.03 - 5.03 (m, 2H), 4.24 - 4.13 (m, 1H), 4.09 - 3.99 (m, 1H), 3.19 - 2.92 (m, 2H), 2.37 - 1.95 (m, 2H), 1.89 - 1.75 (m, 1H), 1.71 - 1.48 (m, 2H), 1.47 - 1.33 (m, 3H), 0.86 (d, *J* = 2.1 Hz, 6H). HRMS m/z: [M+H]^+^ Calculated for C_21_H_25_D_5_N_3_O_5_: 409.2499, Found: 409.2528, m/z: [M+Na]^+^ Calculated for C_21_H_24_D_5_N_3_NaO_5_: 431.2319, Found: 431.2348.

#### Compound *7*

Sodium (2S)-1-hydroxy-2-((S)-4-methyl-2-((((phenyl-d5)methoxy)carbonyl)amino)pentanamido)-3-((S)-2-oxopyrrolidin-3-yl)propane-1-sulfonate (***7***). Yield (68%), mp 125-128 ^0^C. ^1^H NMR (400 MHz, DMSO-d_6_) δ 7.64 (d, *J* = 9.1 Hz, 1H), 7.59 (d, *J* = 9.3 Hz, 1H), 7.49 (d, *J* = 8.1 Hz, 1H), 5.44 (d, *J* = 6.3 Hz, 2H), 5.29 (d, *J* = 6.0 Hz, 1H), 5.05 - 4.98 (m, 2H), 4.03 - 3.87 (m, 2H), 3.17 - 2.95 (m, 2H), 2.24 - 2.02 (m, 2H), 1.83 - 1.50 (m, 3H), 1.49 - 1.37 (m, 2H), 0.84 (d, *J* = 3.0 Hz, 6H). HRMS m/z: [M+H]^+^ Calculated for C_21_H_26_D_5_N_3_NaO_8_S: 513.2043, Found: 513.2054, m/z: [M+Na]^+^ Calculated for C_21_H_25_D_5_N_3_Na_2_O_8_S: 535.1863, Found: 535.1904.

#### Compound *8*

Sodium (5S,8S)-5-isobutyl-3,6,11-trioxo-8-(((S)-2-oxopyrrolidin-3-yl)methyl)-1-(phenyl-d5)-2,10-dioxa-4,7-diazahexadecane-9-sulfonate (***8***). Yield (67%), mp 110-112 ^0^C. ^1^H NMR (400 MHz, DMSO-d_6_) δ 7.87 (d, *J* = 9.3 Hz, 1H), 7.68 (d, *J* = 9.9 Hz, 1H), 7.50 (d, *J* = 9.5 Hz, 1H), 5.11 (d, *J* = 8.9 Hz, 1H), 5.04 - 5.01 (m, 2H), 4.38 - 4.04 (m, 1H), 4.04 - 3.80 (m, 1H), 3.21 - 2.89 (m, 2H), 2.37 - 2.06 (m, 2H), 2.01 (t, *J* = 7.4 Hz, 3H), 1.98 - 1.67 (m, 0H), 1.67 - 1.55 (m, 4H), 1.45 (p, *J* = 7.4 Hz, 2H), 1.39 - 1.33 (m, 1H), 1.32 - 1.16 (m, 4H), 0.93 - 0.76 (m, 9H). HRMS m/z: [M+H]^+^ Calculated for C_27_H_36_D_5_N_3_NaO_9_S: 611.2775, Found: 611.2816.

#### Compound *9*

(Phenyl-d5)methyl-d2 ((S)-4-methyl-1-oxo-1-(((S)-1-oxo-3-((S)-2-oxopyrrolidin-3-yl)propan-2-yl)amino)pentan-2-yl)carbamate (***9***). Yield (70%), mp 45-48 ^0^C. ^1^H NMR (400 MHz, DMSO-d_6_) δ 9.40 (s, 1H), 8.48 (d, *J* = 7.6 Hz, 1H), 7.63 (s, 1H), 7.47 (d, *J* = 8.3 Hz, 1H), 4.23 - 4.17 (m, 1H), 4.12 - 4.03 (m, 1H), 3.23 - 2.95 (m, 2H), 2.37 - 2.03 (m, 3H), 1.93 - 1.81 (m, 1H), 1.74 - 1.56 (m, 3H), 1.56 - 1.36 (m, 1H), 0.87 (d, *J* = 6.6 Hz, 6H). HRMS m/z: [M+H]^+^ Calculated for C_21_H_23_D_7_N_3_O_5_: 411.2625, Found: 411.2622, m/z: [M+Na]^+^ Calculated for C_21_H_22_D_7_N_3_NaO_5_: 433.2445, Found: 433.2440.

#### Compound *10*

Sodium (2S)-1-hydroxy-2-((S)-4-methyl-2-((((phenyl-d5)methoxy-d2)carbonyl)amino)pentanamido)-3-((S)-2-oxopyrrolidin-3-yl)propane-1-sulfonate (***10***). Yield (82%), mp 120-122 ^0^C. ^1^H NMR (400 MHz, DMSO-d_6_) δ 7.66 (d, *J* = 9.1 Hz, 0H), 7.60 (d, *J* = 9.5 Hz, 1H), 7.50 (s, 1H), 5.51 (d, *J* = 6.2 Hz, 1H), 5.34 (d, *J* = 6.0 Hz, 0H), 4.06 - 3.97 (m, 1H), 3.97 - 3.90 (m, 1H), 3.15 - 3.07 (m, 2H), 2.21 - 2.01 (m, 3H), 2.01 - 1.71 (m, 1H), 1.71 - 1.49 (m, 4H), 1.49 - 1.31 (m, 2H), 0.87 (d, *J* = 4.7 Hz, 6H). HRMS m/z: [M+H]^+^ Calculated for C_21_H_24_D_7_N_3_NaO_8_S: 515.2169, Found: 515.2152.

#### Compound *11*

Sodium (5S,8S)-5-isobutyl-3,6,11-trioxo-8-(((S)-2-oxopyrrolidin-3-yl)methyl)-1-(phenyl-d5)-2,10-dioxa-4,7-diazahexadecane-9-sulfonate-1, 1-d2 (***11***). Yield (44%), mp 100-105 ^0^C. ^1^H NMR (400 MHz, DMSO-d_6_) δ 7.64 (d, *J* = 8.7 Hz, 1H), 7.48 (s, 1H), 7.35 (d, *J* = 8.0 Hz, 1H), 5.13 (d, *J* = 8.8 Hz, 1H), 4.16 - 4.03 (m, 1H), 4.05 - 3.87 (m, 1H), 3.21 - 2.87 (m, 2H), 2.40 - 1.99 (m, 4H), 1.99 - 1.76 (m, 1H), 1.76 - 1.34 (m, 7H), 1.34 - 1.18 (m, 4H), 0.96 - 0.69 (m, 9H). HRMS m/z: [M+H]^+^ Calculated for C_27_H_34_D_7_N_3_NaO_9_S: 613.2900, Found: 613.2899, m/z: [M]^-^ Calculated for C_27_H_33_D_7_N_3_O_9_S: 589.2924, Found: 589.2940.

**Figure S1.**
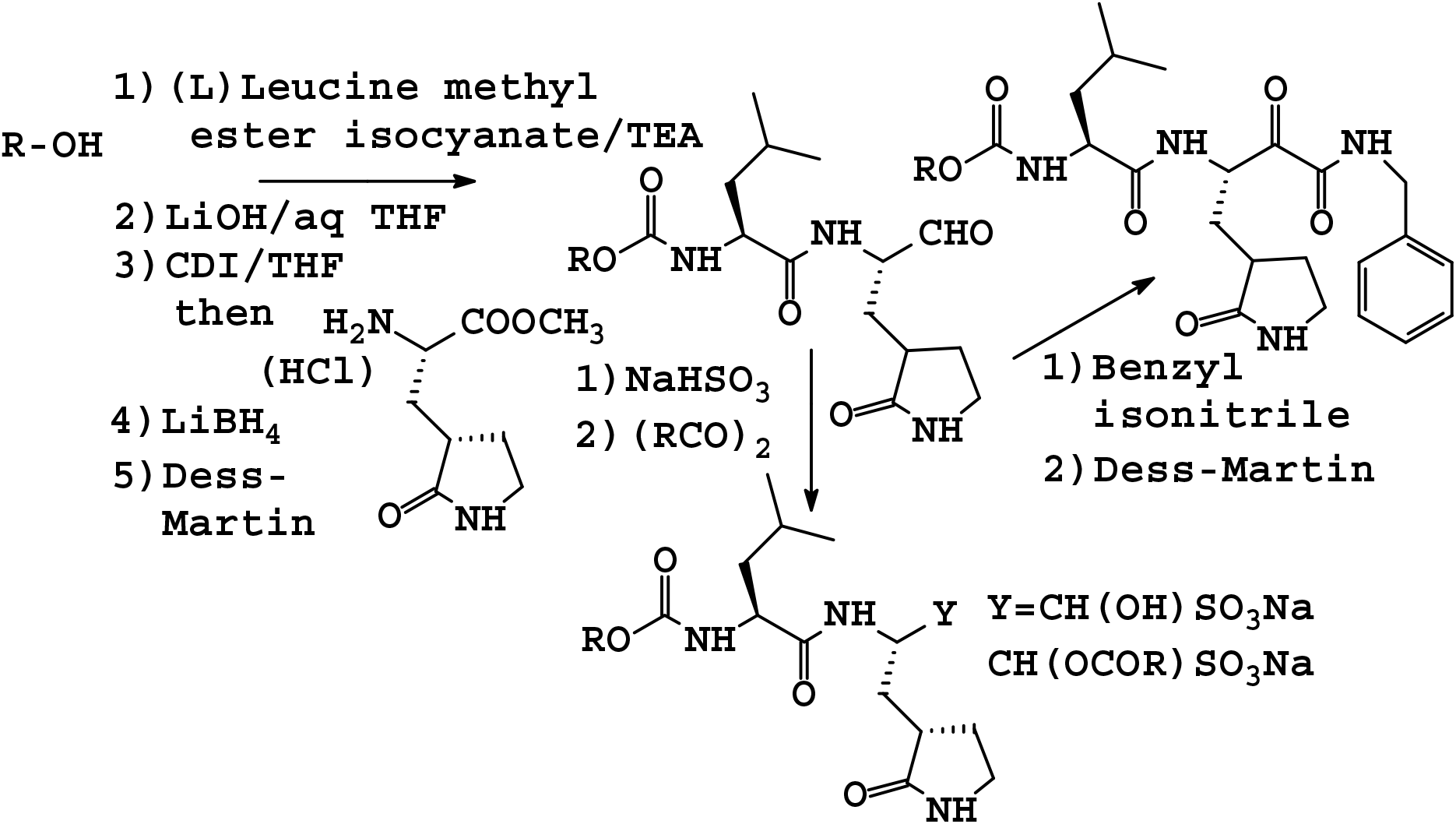
Synthesis of deuterated inhibitors ***1-11***

**Figure S2.**
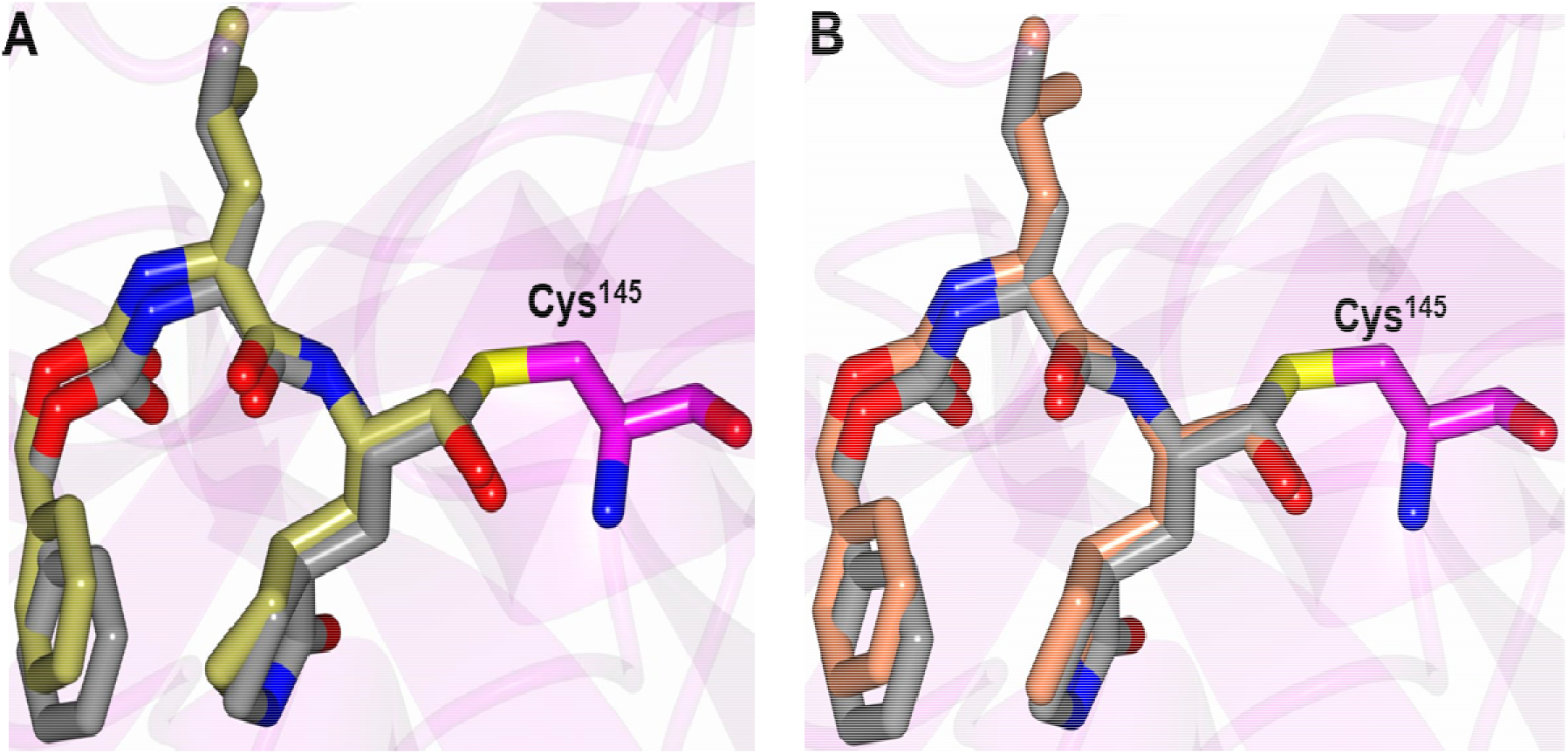
Structure of SARS-CoV-2 3CLpro in complex with compound ***2*** (gray) superimposed with (A) SARS-CoV-2 3CLpro with GC376 (PDB 6WTJ, gold) and (B) SARS-CoV-2 3CLpro with GC373 (PDB 6WTK, coral).

**Figure S3.**
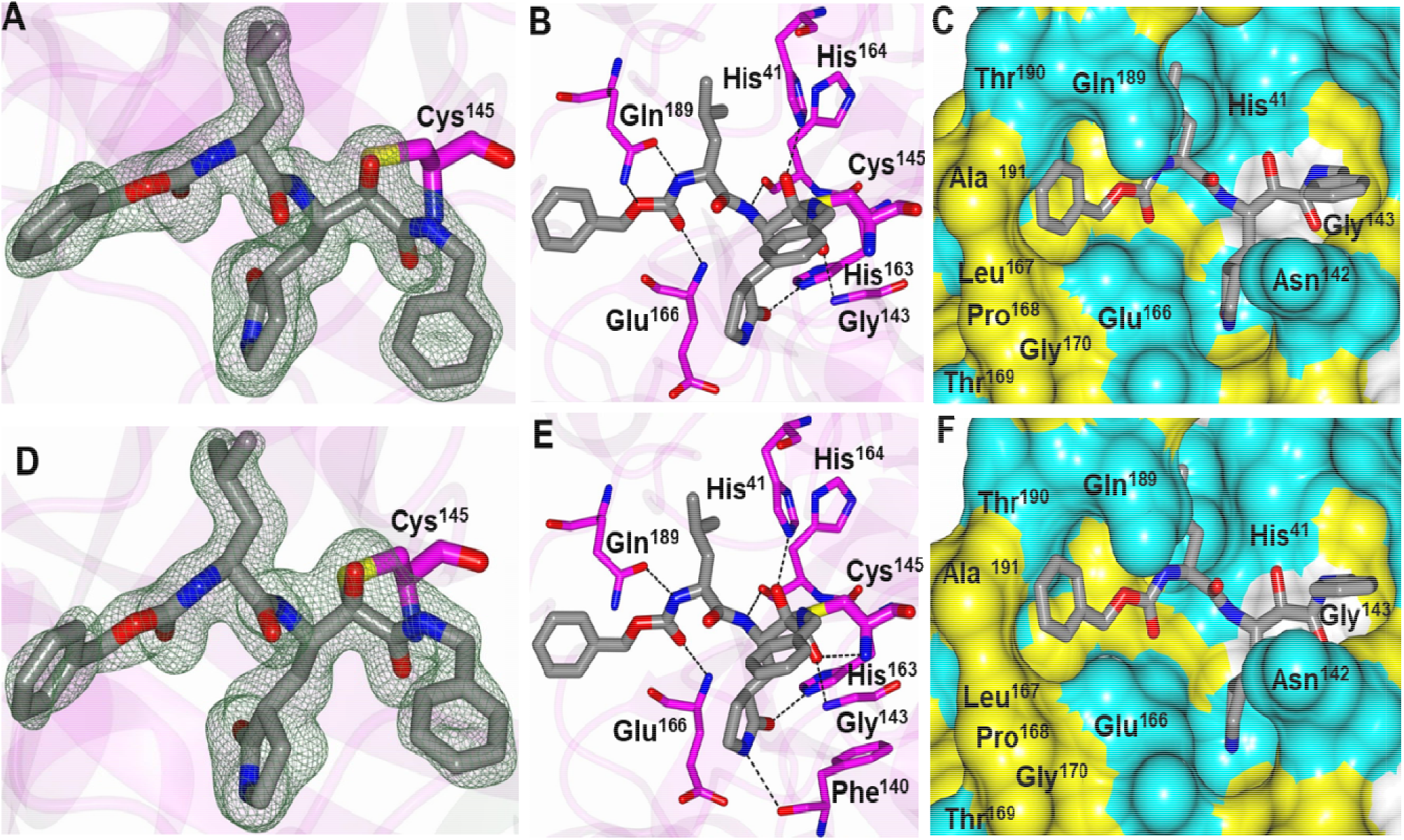
Cocrystal structures of SARS-CoV 3CLpro (A, B, C) and SARS-CoV-2 3CLpro (D, E, F) in complex with compound ***5***. Panels A and D show *F_o_-F_c_* omit maps (green mesh) contoured at 3σ. Panels B and E show hydrogen bond interactions (dashed lines) between the inhibitor and the 3CL protease. Panels C and F show electrostatic surface representation of the binding pocket occupied by the inhibitor. Neighboring residues are colored yellow (nonpolar), cyan (polar), and white (weakly polar).

**Figure S4.**
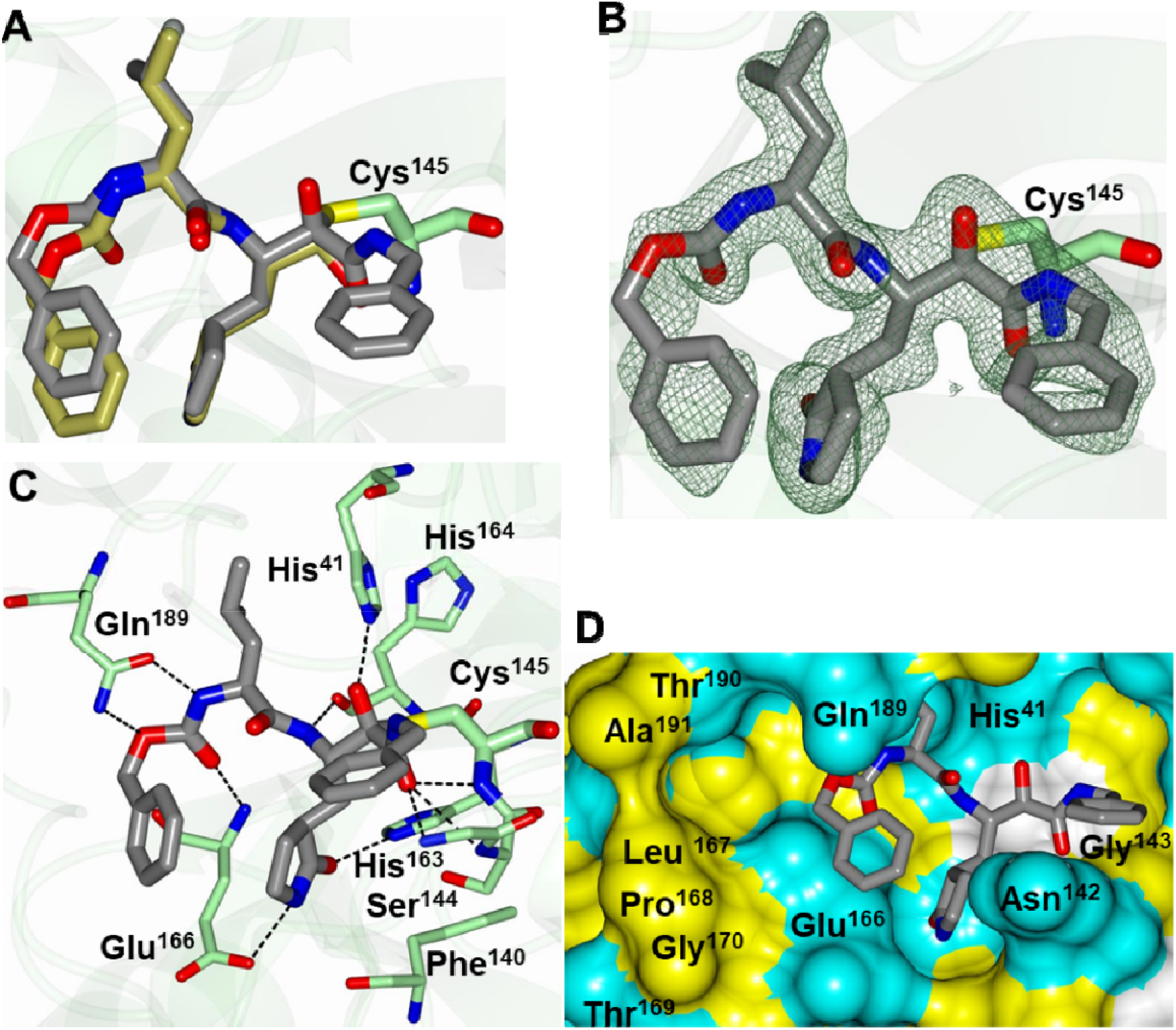
Structure of SARS-CoV-2 3CLpro in complex with compound ***5*** associated with subunit B. (A) Structure of compound **2** (gold) superimposed onto compound **5** in subunit B (gray) showing the similar binding modes. (B) *F_o_-F_c_* omit map (green mesh) contoured at 3σ. (C) Hydrogen bond interactions (dashed lines) between the inhibitor and the 3CL protease. (D) Electrostatic surface representation of the binding pocket occupied by the inhibitor. Neighboring residues are colored yellow (nonpolar), cyan (polar), and white (weakly polar).

**Table S1.**
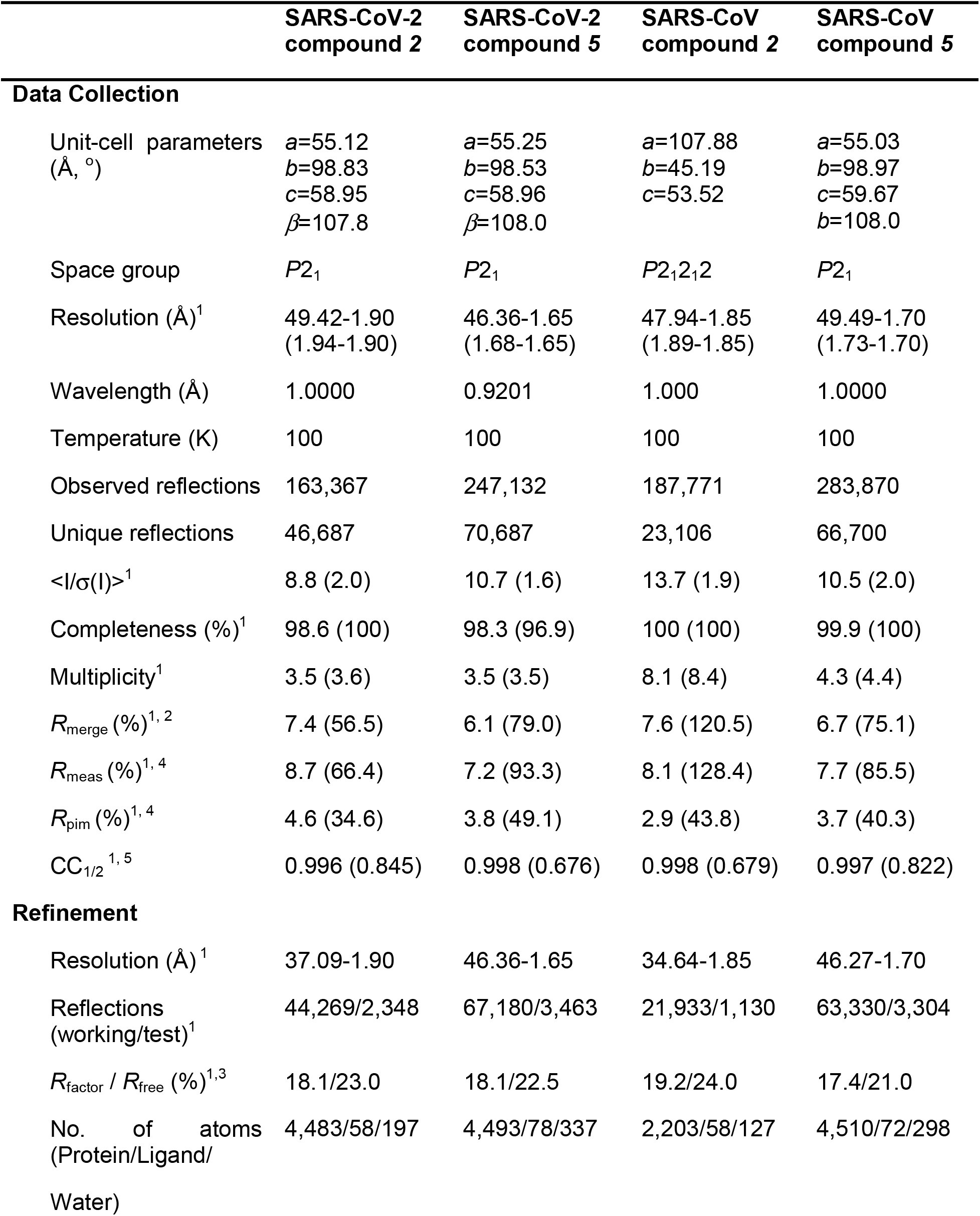
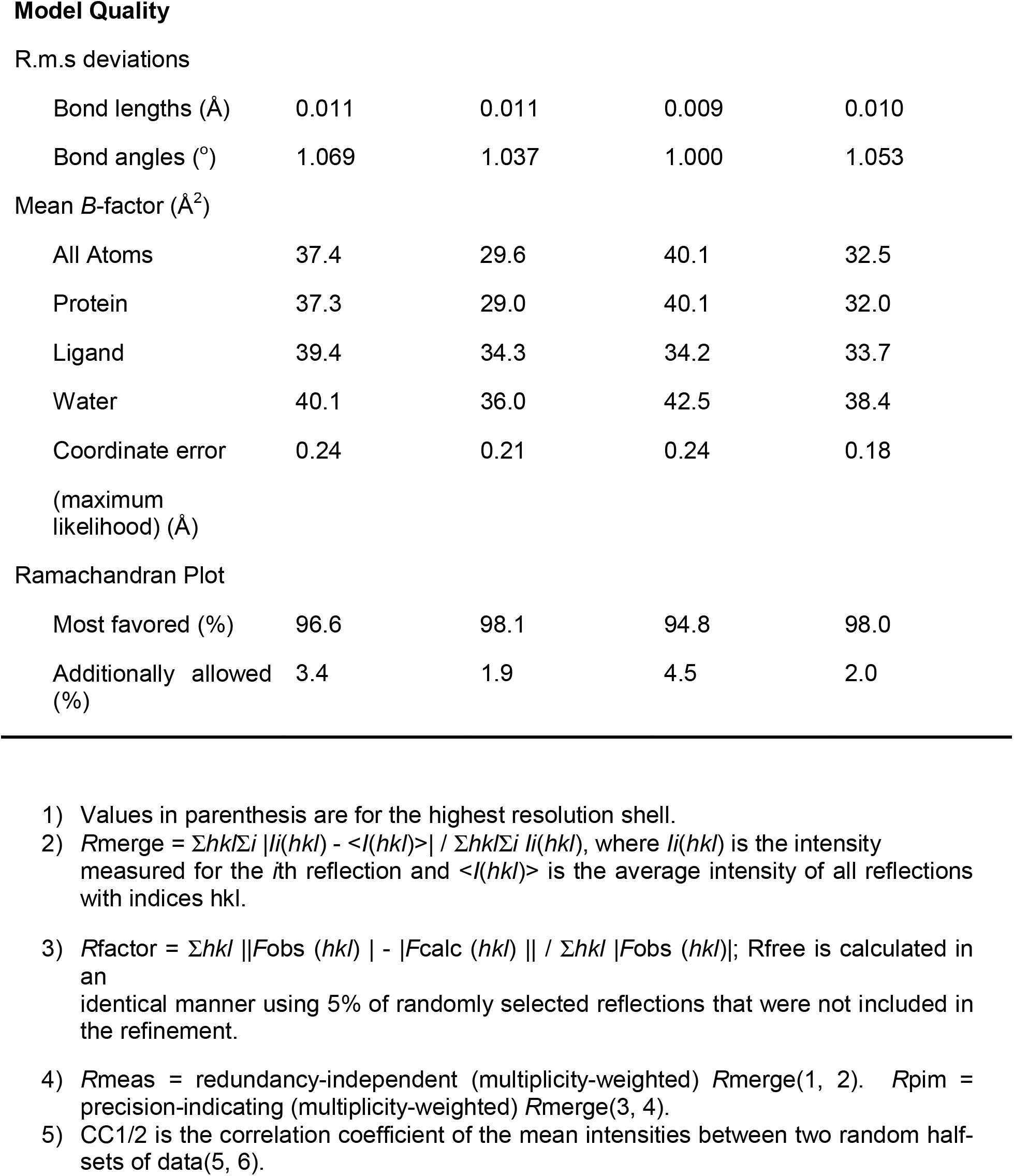
Crystallographic data for SARS-CoV and SARS-CoV-2 3CLpro structures.

